# ZMS-6659, a selective Polθ helicase inhibitor, demonstrates anti-tumor activity as monotherapy and synergy effect with PARP inhibitors and topoisomerase inhibitors

**DOI:** 10.1101/2025.04.19.649634

**Authors:** Lei Jiang, Jiajia Li, Lu Liu, Mengying Li, Wenjing Li, Zhen Li, Shuo Qian, Mingyue Shi, Jing Dai, Chunxia Ao, Ying Qu, Zhengtao Li, Li Zhou, Chen Yang, Xiyuan Wang, Renhong Tang, Feng Zhou, Gang Liu, Liting Xue

**Affiliations:** State Key Laboratory of Neurology and Oncology Drug Development, Nanjing 210023, P.R. China; Simcere Zaiming Pharmaceutical Co, Ltd., Shanghai 200000, P.R. China; School of Pharmaceutical Sciences, Tsinghua University, Beijing 100084, P.R. China; Department of Gynecologic Oncology, Fudan University Shanghai Cancer Center, Shanghai, 200032, P.R. China; School of Science, China Pharmaceutical University, Nanjing 211198, P.R. China

**Keywords:** Polθ inhibitor, helicase domain, PARP inhibitor, MMEJ, synthetic lethality

## Abstract

The repair of DNA double strand break (DSB) is crucial for genome stability. Homologous recombination (HR) is a high-fidelity, error-free DSB repair pathway, and its dysfunction leads to genome instability and can result in tumorigenesis. In HR deficient contexts, microhomology-mediated end joining (MMEJ), dependent on DNA polymerase theta (Polθ), is up-regulated to serve as a backup pathway for DSB repair. Several studies have proved that Polθ inhibition causes synthetic lethality with HR deficiency, and Polθ emerges as a potential DNA damage repair (DDR) target for the treatment of HR deficient tumors both as single agent or in combination with PARP inhibitors (PARPi). In addition, evidences show that the blockage of Polθ can enhance the sensitivity to DNA damage agents. Herein, we characterized ZMS-6659, a highly potent small-molecule Polθ inhibitor targeting the helicase domain. ZMS-6659 induces synthetic lethality in HR deficient models, elicits synergistic effects with even low dose of PARPi and delays PARPi resistance without additional hematotoxicity. Besides, ZMS-6659 also enhance the sensitivity to topoisomerase inhibitors in HR proficient cell lines. These findings support the clinical exploration of ZMS-6659 in combination with PARPi in HR deficient tumors, and suggest a potential application expansion beyond HR deficiency.

## Introduction

The repair of DNA double-strand break (DSB) is crucial for genome stability and cell survival. There are three main DSB repair pathways: homologous recombination (HR), non-homologous end joining (NHEJ), and microhomology-mediated end joining (MMEJ). Among them, HR is a high-fidelity, error-free DSB repair pathway which is crucial for preserving genome stability, while NHEJ and MMEJ are error-prone pathways and may result in mutations at DSB repair sites^1^. The dysfunction of HR confers cell genome instability and leads to tumorigenesis, which is prevalent in 10% gynecologic tumors, especially frequently occurring in approximately 20%-30% ovarian cancer and breast cancer^2^.

In situation of HR deficiency, MMEJ pathway is upregulated to compensate for defective DSB repair^3^. This error-prone repair mechanism critically depends on DNA polymerase theta (Polθ), a bifunctional enzyme comprising an N-terminal superfamily 2 HEL308-type helicase domain and a C-terminal A-family DNA polymerase domain^4^. The helicase domain mediates the displacement of replication protein A (RPA) from single-stranded DNA (ssDNA) and ssDNA annealing through microhomology recognition. The polymerase domain executes strand extension and gap-filling synthesis using microhomology-aligned templates. These coordinated enzymatic activities enable Polθ to complete the MMEJ repair process.

The clinical application of PARP inhibitor (PARPi) is the first clinically validated synthetic lethality through interaction with HR deficiency^5^. However, emerging evidence reveals Polθ-mediated MMEJ as a dual-edged sword, functioning both as an exploitable vulnerability for HR deficiency and as a PARPi resistance mechanism through backup DNA repair due to its nature of error-prone repair^6,7^. While first-generation Polθ inhibitors targeting the polymerase domain, such as ART558 and RP-6685, have demonstrated preclinical synergy with PARPi, the weak cellular potency at hundred nanomolar even micromolar level limits their translational potential and is insufficient to fully evaluate the safety and efficacy of targeting Polθ in clinical application^8,9^.

Preclinical models show that Polθ knockout enhances tumor sensitivity to diverse genotoxic agents, including chemotherapy and radiotherapy^10–14^. These evidences create rationale for exploring Polθ inhibition in combination with topoisomerase inhibitors (TOPOi), which induce lethal DNA double-strand breaks (DSBs), and are both widely used as chemotherapy agents and serve as the precision medicine hot spot antibody-drug conjugate (ADC) payloads in clinical practice^15^.

Herein, we characterized ZMS-6659, a small-molecule Polθ inhibitor eliciting highly potent biochemical and cellular activity through selective targeting of the Polθ helicase domain. ZMS-6659 not only induces synthetic lethality in HR deficient models but also exhibits synergistic effects with PARPi and delays PARPi resistance development. Notably, these effects occur without additional hematotoxicity profiles. Furthermore, the therapeutic potential of ZMS-6659 in combination with TOPOi in HR proficient cell lines was demonstrated. These findings collectively support the clinical exploration of ZMS-6659 in combination with PARPi in HR deficient tumors, and suggest a potential application expansion beyond HR deficiency.

## Results

### ZMS-6659 is a highly potent specific Polθ helicase inhibitor

The Polθ protein consists of an N-terminal helicase domain (superfamily 2 HEL308-type) and a C-terminal DNA polymerase domain (A-family) (Figure 1A). Several Polθ inhibitors targeting the polymerase domain, such as ART558, have been reported and displayed potencies at micromolar level^8,9,17^. Here, we developed a novel small molecular Polθ helicase inhibitor ZMS-6659. It inhibited Polθ helicase activity with an IC_50_ of 16.80±1.28 nM (Figure 1B) and did not inhibit Polθ polymerase activity (IC_50_>6000 nM) (Figure 1C). Both novobiocin and ART558 displayed moderate biochemical activity with IC_50_ of 75046.71±4726.15 nM and 40.65 nM respectively. To gain insight into the binding mode of ZMS-6659, a kinetic assay was conducted. The result showed that the V_max_ remained unchanged while the K_m_ increased along with the elevating of ZMS-6659 concentration, revealing that ZMS-6659 is an ATP-competitive Polθ helicase inhibitor (Figure 1D & Supplementary Table 1). Next, we investigated the selectivity of ZMS-6659 against several helicases which also harbor SF2 DNA helicase domains^18^. ZMS-6659 showed no significant inhibitory activity against these helicases (IC_50_>10000 nM) except for WRN, of which ZMS-6659 only had a weak inhibitory effect with an IC_50_ of 8040.31 nM, demonstrating good selectivity (Figure 1E).

**Figure 1.**
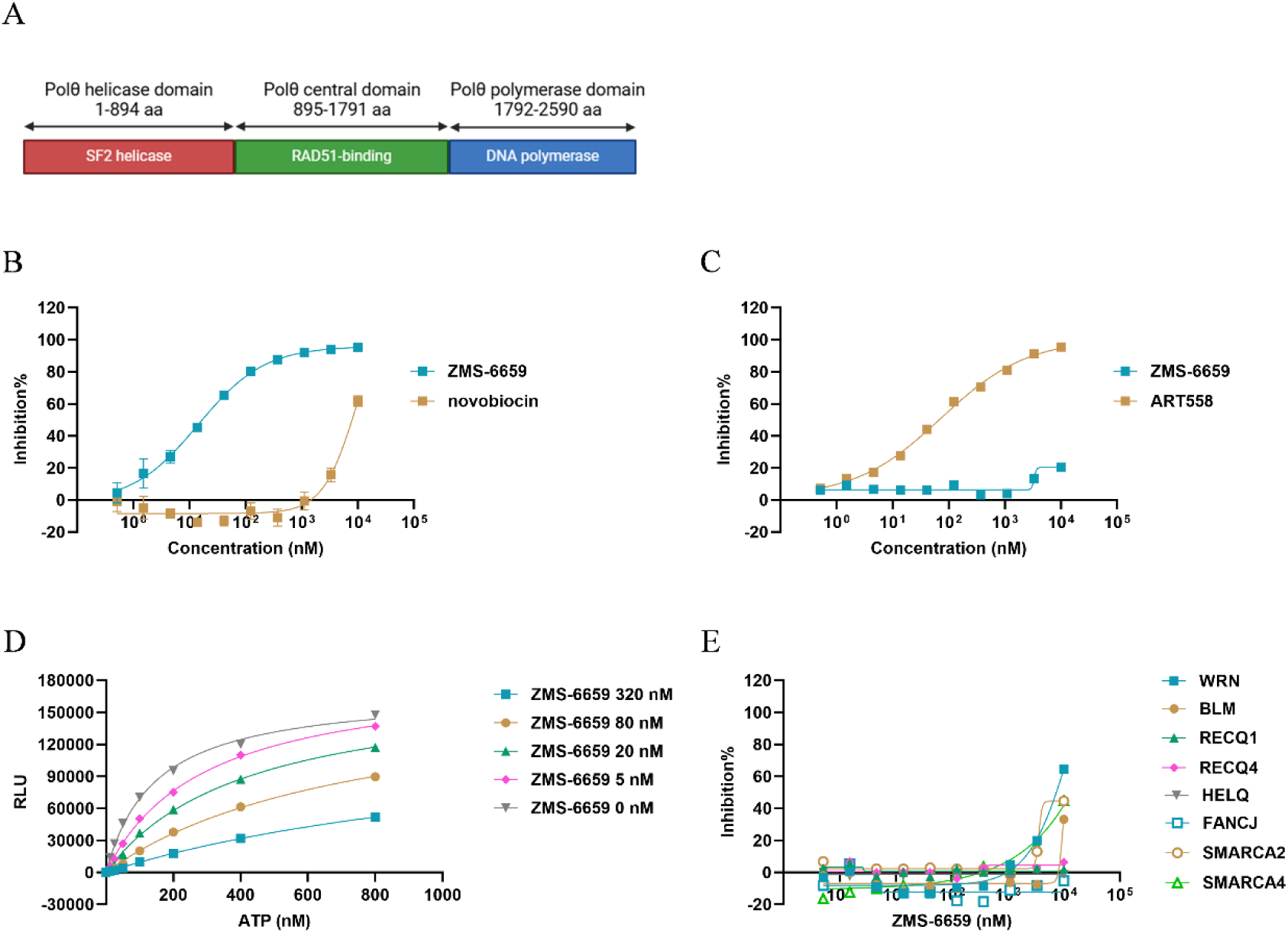
ZMS-6659 is a highly potent specific Polθ helicase inhibitor. (A) Schematic representation of Polθ. (B) The Polθ helicase inhibition activity of ZMS-6659. Novobiocin was served as a positive control. Each data point represents the mean ± SD from 6 biological replicates of 3 individual experiments (n=6). (C) The Polθ polymerase inhibition activity of ZMS-6659. ART-558 was served as a positive control. (D) The ATP competition assay. (E) The SF2 DNA helicase inhibition activity of ZMS-6659. (C-E) Each data point represents the mean from 2 biological replicates (n=2).

### ZMS-6659 causes synthetic lethality with HR deficiency through the inhibition of MMEJ pathway

Polθ plays an essential role in the MMEJ pathway, which serves as a backup pathway for HR in DSB repair and is upregulated when HR function is defective^3,6^. Series of studies have demonstrated that the inhibition of Polθ polymerase domain confers synthetic lethality with HR deficiency^8,9^. To verify whether targeting helicase domain of Polθ has similar functions, we first evaluated the ability of ZMS-6659 to inhibit Polθ-mediated MMEJ repair in the pre-described reporter assay. The result demonstrated that ZMS-6659 robustly inhibited the cellular MMEJ axis with an IC_50_ of 1.69±0.67 nM, which is much stronger than the reported Polθ polymerase inhibitor ART558 (Figure 2A). We also conducted a 7-days cell proliferation assay in isogenic DLD-1 cells with or without a truncating mutation in BRCA2 (DLD-1 BRCA2-/- and DLD-1 parental cells) and a number of non-malignant cells. ZMS-6659 strongly inhibited the proliferation of HR deficient DLD-1 BRCA2-/- cells with an IC_50_ of 10.69 ±6.96 nM, and showed a >900× selectivity folds over DLD-1 parental cells as well as non-malignant cells (Figure 2B & 2C). Accordingly, phosphorylated H2A (γH2AX), a DNA damage related biomarker, was upregulated in a dose-dependent manner in DLD-1 BRCA2-/- cells (Figure 2D). These results demonstrated that ZMS-6659 elicited synthetic lethality with HR deficiency *in vitro* and had good selectivity for HR proficient cells.

**Figure 2.**
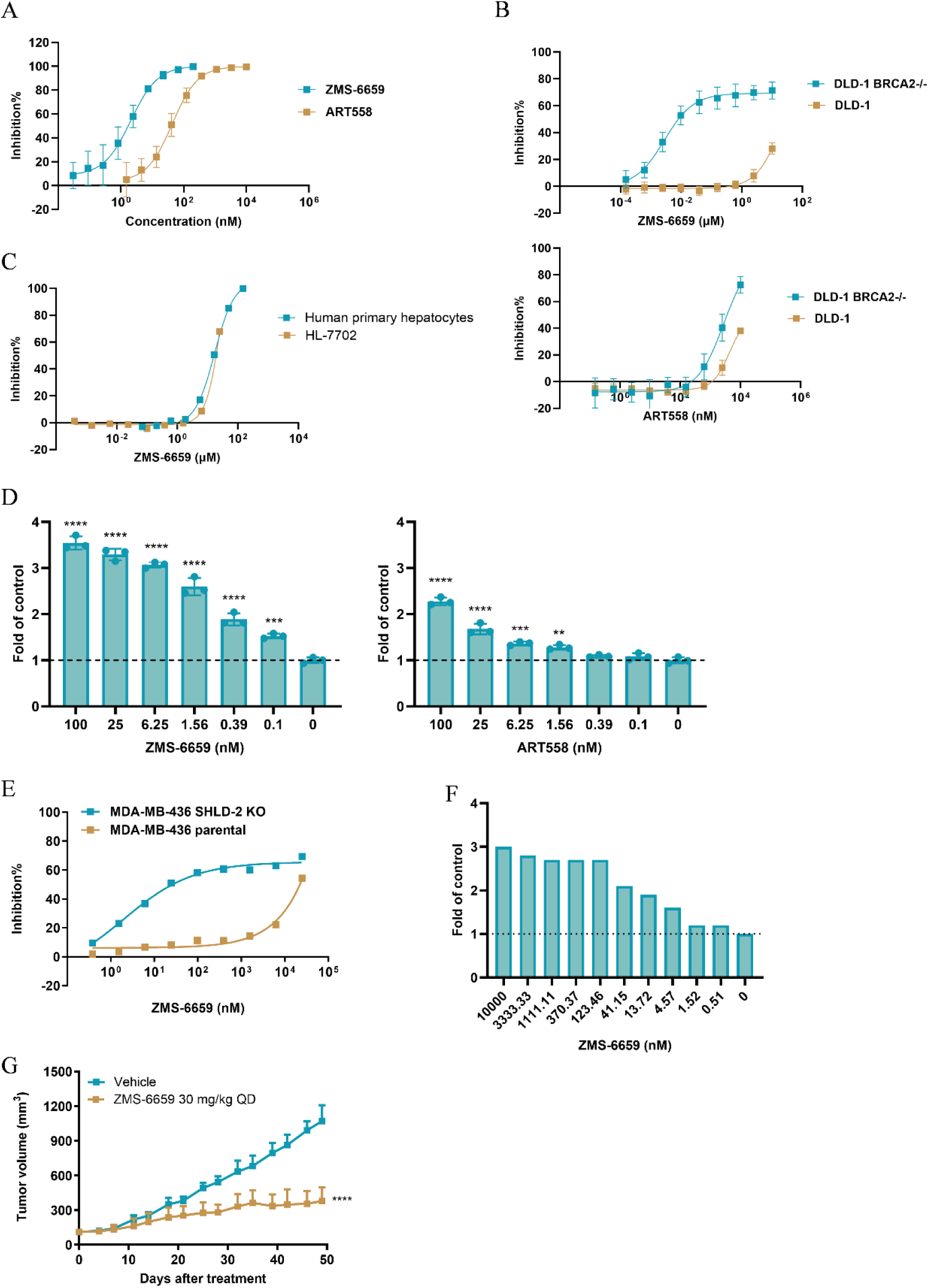
ZMS-6659 induces synthetic lethality with HR deficiency and Shieldin complex deficiency further enhances the activity of ZMS-6659. (A) The MMEJ pathway inhibition activity of ZMS-6659. ART-558 was served as a positive control. Each data point represents the mean ± SD from 6 biological replicates of 3 individual experiments (n=6). (B) Cell proliferation inhibition activities of ZMS-6659 or ART558 in DLD-1 BRCA2-/- and DLD1 parental cells. Each data point represents the mean ± SD from 6 biological replicates of 3 individual experiments (n=6). (C) Cell proliferation inhibition activities of ZMS-6659 in HL-7702 cells and human primary hepatocytes. Each data point represents the mean from 2 biological replicates (n=2) in HL-7702 cells and 3 biological replicates (n=3) in human primary hepatocytes. (D) γH2AX accumulation in DLD-1 BRCA2-/- cells after the treatment of indicated concentrations of ZMS-6659 or ART558 for 72 h. Each bar represents the mean ± SD from 3 biological replicates (n=3). Statistical significance relative to vehicle was established by unpaired t test. ** p<0.01, *** p<0.001, ****p<0.0001. (E) Cell proliferation inhibition activities of ZMS-6659 in MDA-MB-436 SHLD2-/- and MDA-MB-436 parental cells. Each data point represents the mean from 2 biological replicates (n=2). (F) γH2AX accumulation in MDA-MB-436 SHLD2-/- cells after the treatment of indicated concentrations of ZMS-6659 for 72 h. (G) Tumor growth inhibition of ZMS-6659 in MDA-MB-436 SHLD2-/- xenograft mice model. Animals were treated for 49 days with either vehicle (n = 6) or ZMS-6659 (30 mg/kg, QD, n=6). Statistical significance relative to vehicle was established by two-way ANOVA with Tukey’s multiple comparison test. ****p<0.0001.

### The deficiency of Shieldin complex enhances the sensitivity to ZMS-6659

We tested the activity of ZMS-6659 in a natural BRCA1 mutation cell line MDA-MB-436, and observed that ZMS-6659 displayed weak potency in the 7-day cell proliferation assay and required a prolonged 14-days treatment to elicit the strong synergy effect with HR deficiency (Supplementary Figure 1A & 1B). Meanwhile, ZMS-6659 as a single agent did not display significant anti-tumor efficacy in either DLD-1 BRCA2 -/- or MDA-MB-436 xenograft model (data not shown). These findings prompted us to investigate potential compensatory DSB repair mechanisms beyond HR and microhomology-mediated end joining (MMEJ). We focused on NHEJ, a competing DSB repair pathway mediated by the 53BP1-recruited Shieldin complex. Previous studies have demonstrated that depletion of Shieldin components (MAD2L2, SHLD1, SHLD2, SHLD3) enhances sensitivity to Polθ inhibitors in BRCA1-deficient cells^8^. To test this mechanism in our system, we compared ZMS-6659 activity in isogenic BRCA1-mutant MDA-MB-436 cells with or without SHLD2 knockout (MDA-MB-436 SHLD2-/- and MDA-MB-436 parental cells). The MDA-MB-436 SHLD2-/- cells exhibited dramatically increased sensitivity to ZMS-6659 compared to parental cells (Figure 2E), accompanied by significantly elevated γH2AX levels (Figure 2F). ZMS-6659 alone also elicited significant anti-tumor efficacy in MDA-MB-436 SHLD2-/- tumor xenograft model (Figure 2G). These results collectively demonstrate that Shieldin complex deficiency in BRCA1-deficient models potentiates the sensitivity to Polθ inhibition.

### Combination of ZMS-6659 and PARPi produces synergetic anti-tumor effects without additional hematotoxicity

PARPi have exhibited clinical efficacy in HR deficient tumors through synthetic lethality, and emerging evidences suggest that knockout of Polθ may enhance PARPi sensitivity^8,19^. Hence, we evaluated the combination of ZMS-6659 with olaparib in DLD-1 BRCA2-/- cells. The combination demonstrated synergistic anti-proliferative effects accompanied by elevated γH2AX levels (Figure 3A & 3B), indicating the enhanced DNA damage accumulation. This synergistic effect was also observed in multiple HR-deficient cell lines, including MDA-MB-436 and Capan-1 cells (Figure 3C & 3D), and extended to other PARP inhibitors including niraparib, talazoparib, and the PARP1-selective inhibitor AZD5305 (Supplementary Figure 1C-1E). Furthermore, we validated this combinatorial efficacy in a clinically relevant *ex vivo* model using HR-deficient ovarian cancer patient-derived tumor tissue, demonstrating translational potential of this therapeutic approach (Figure 3E).

**Figure 3.**
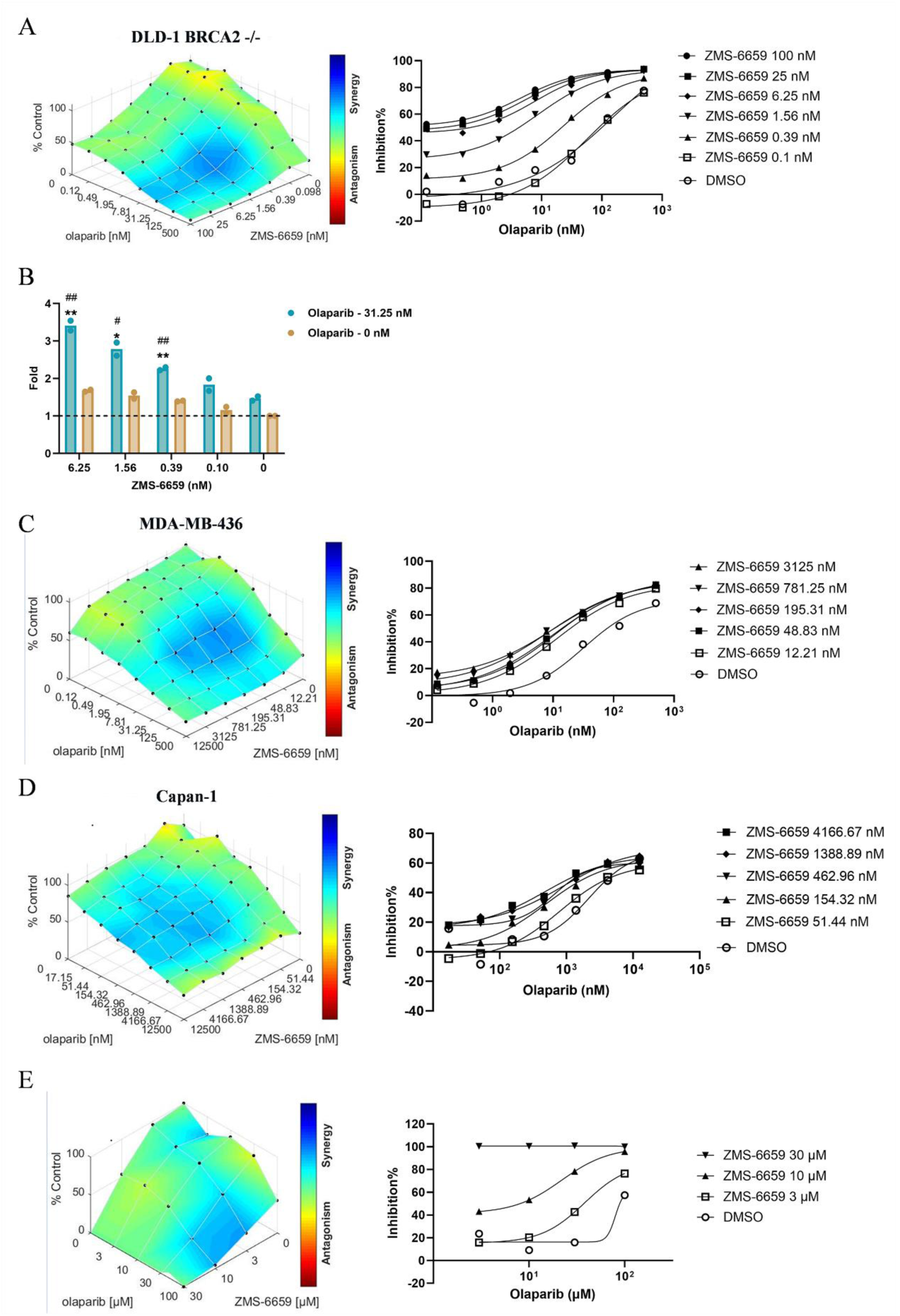
ZMS-6659 elicits synergy effect with PARPi and delays the resistance to PARPi. (A) The combination of ZMS-6659 and olaparib in DLD-1 BRCA2-/- cells. Each data point represents the mean from 2 biological replicates (n=2). (B) γH2AX accumulation in DLD-1 BRCA2-/- cells after the combination treatment of ZMS-6659 and olaparib. Each bar represents the mean from 2 biological replicates (n=2). Statistical significance was established by unpaired t test. *p<0.05, **p<0.01 versus olaparib. # p<0.05, ## p<0.01 versus ZMS-6659 at corresponding concentration. (C) The combination of ZMS-6659 and olaparib in MDA-MB-436 cells. Each data point represents the mean from 2 biological replicates (n=2). (D) The combination of ZMS-6659 and olaparib in Capan-1 cells. Each data point represents the mean from 2 biological replicates (n=2). (E) The combination of ZMS-6659 and olaparib in a 3D-bioprinted clinically relevant HR-deficient ovarian cancer patient-derived *ex vivo* model. Each data point represents the mean from 3 biological replicates (n=3).

To validate these findings *in vivo*, we conducted a multiple-dose evaluation in the DLD-1 BRCA2-/- xenograft model. Mice received daily oral administration of ZMS-6659 (0.3, 1, or 10 mg/kg) in combination with 50 mg/kg olaparib, or 10 mg/kg ZMS-6659 in combination with lower dose of olaparib (15 mg/kg). The combination of 0.3 mg/kg ZMS-6659 and 50 mg/kg olaparib significantly suppressed tumor growth, and the combination of higher doses (1 or 10 mg/kg) of ZMS-6659 and 50 mg/kg olaparib further induced tumor regression without weight loss (Figure 4A & Supplementary Figure S2), demonstrating superior efficacy over monotherapy. Notably, the combination of 10 mg/kg ZMS-6659 with 15 mg/kg olaparib maintained therapeutic superiority over olaparib alone (Figure 4B), suggesting preserved synergy at lower PARPi exposure levels. These findings supported the potential for reducing PARPi exposure and toxicity while maintaining efficacy in clinical applications. To correlate the anti-tumor efficacy with pharmacokinetic (PK), plasma samples were collected at different timepoint after the last dosing. The result revealed that the plasma concentrations of ZMS-6659 at 1 mg/kg maintained above the cellular MMEJ IC_50-total_ during the dosing interval (Figure 4C), suggesting that high exposure to maintain sustained MMEJ inhibition is required for achieving maximum efficacy. In a subsequent study, we conducted a comparative analysis of efficacy between combination therapy and high-dose olaparib monotherapy. Mice received daily oral administration of ZMS-6659 (1 mg/kg) combined with 50 mg/kg olaparib, or high dose of olaparib (200 mg/kg) alone. The ZMS-6659/olaparib combination demonstrated superior anti-tumor efficacy compared to olaparib alone (Figure 4D). Notably, PK analysis further indicated that the AUC_0-24_ of olaparib at 200 mg/kg reached 53 μg·h/mL, and its unbound drug exposure was comparable to that of 300 mg olaparib tablet in human. Collectively, these findings reinforce the rationale for clinical exploration of ZMS-6659 in combination with PARPi.

**Figure 4.**
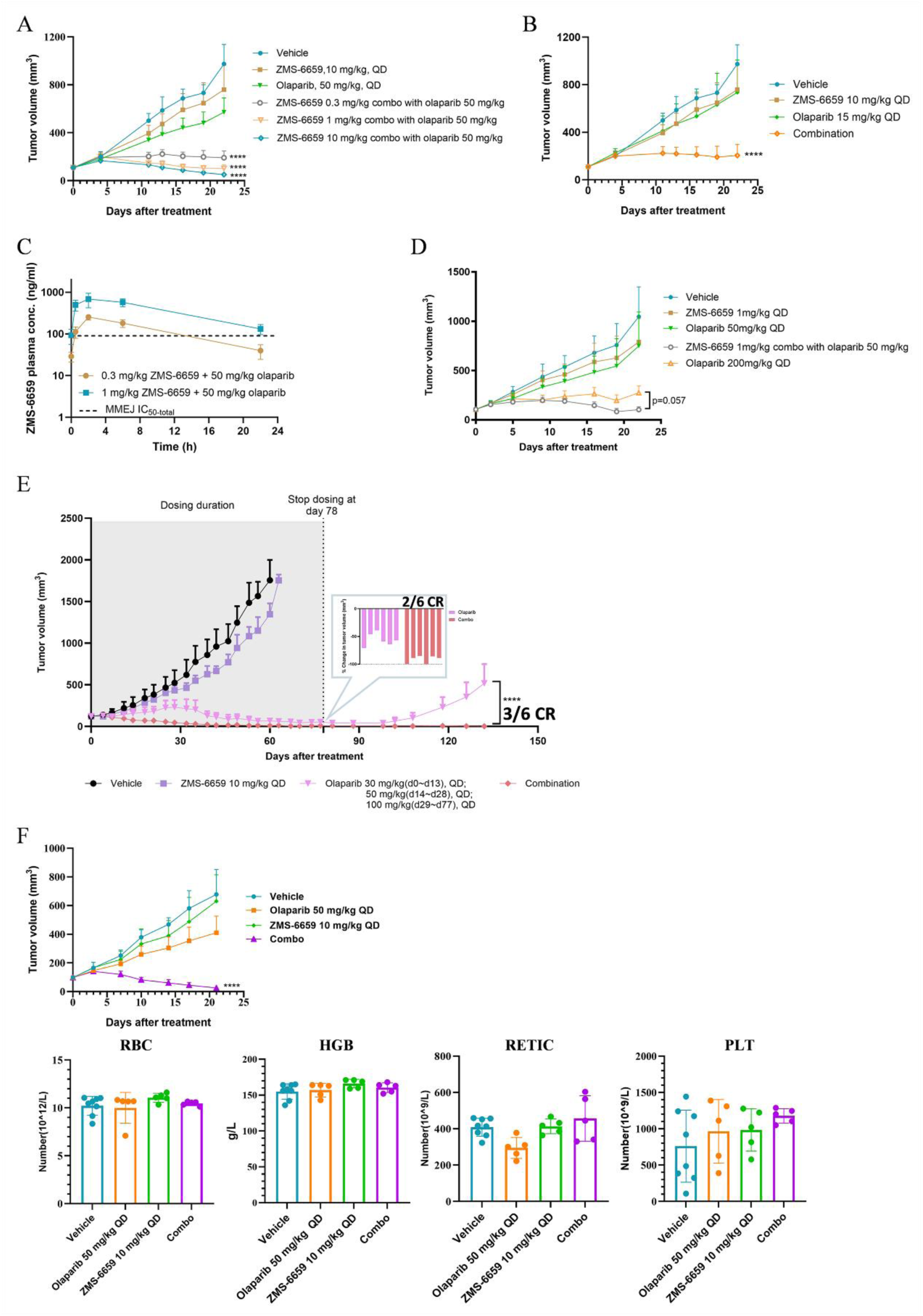
ZMS-6659 and PARPi combination produces synergetic anti-tumor effects without additional hematotoxicity. (A) Tumor growth inhibition of ZMS-6659 and olaparib combination treatment in DLD-1 BRCA2 -/- xenograft mice model. Animals were treated for 22 days with either vehicle (n=8), or ZMS-6659 (10 mg/kg, QD, n=8), or olaparib (50 mg/kg, QD, n=8), or ZMS-6659 (0.3/1/10 mg/kg, QD) and olaparib (50 mg/kg, QD) combination (n=8). ****p<0.0001 versus olaparib (50 mg/kg, QD). (B) Tumor growth inhibition of ZMS-6659 and olaparib combination treatment in DLD-1 BRCA2-/- xenograft mice model. Animals were treated for 22 days with either vehicle (n=8), or ZMS-6659 (10 mg/kg, QD, n=8), or olaparib (15 mg/kg, QD, n=8), or ZMS-6659 (10 mg/kg, QD) and olaparib (15 mg/kg, QD) combination (n=8). ****p<0.0001 versus olaparib (15 mg/kg, QD). (C) Plasma concentrations of ZMS-6659. Blood samples from the experiment of Fig 3E were collected at indicated time points after the final dosing. Each data point represents the mean ± SD from 4 biological replicates (n=4). The dotted line depicts ZMS-6659 plasma concentration required to cover cellular MMEJ IC_50_, which was calculated by dividing the cellular IC_50_ by the plasma unbound fraction. (D) Tumor growth inhibition of ZMS-6659 and olaparib combination treatment in DLD-1 BRCA2-/- xenograft mice model. Animals were treated for 22 days with either vehicle (n=8), or ZMS-6659 (1 mg/kg, QD, n=8), or olaparib (50/200 mg/kg, QD, n=8), or ZMS-6659 (1 mg/kg, QD) and olaparib (50 mg/kg, QD) combination (n=8). (E) Tumor growth inhibition of ZMS-6659 and olaparib combination treatment in MDA-MB-436 xenograft mice model. Animals were treated with vehicle from day 0 to day 60 (n=4 at day 60, 2 mice were sacrificed at day 60 according to animal welfare policy), or with ZMS-6659 from day 0 to day 63 (10 mg/kg, QD, n=6), or with olaparib for indicated time (30/50/100 mg/kg, QD, n=6), or with combination group from day 0 to day 77 (n=6). ****p<0.0001 versus olaparib alone. CR: complete regression. (F) Tumor growth inhibition (top) and blood sample tests (bottom) of ZMS-6659 and olaparib combination treatment in DLD-1 BRCA2-/- xenograft mice model. Animals were treated for 21 days with either vehicle (n=8), or ZMS-6659 (10 mg/kg, QD, n=8), or olaparib (50 mg/kg, QD, n=8), or ZMS-6659 (10 mg/kg, QD) and olaparib (50 mg/kg, QD) combination (n=8). Blood samples were collected after the final dosing. Each bar represents the mean ± SD from 8 (vehicle, n=8) or 5 (other groups, n=5) biological replicates. (A, B, D, E, F) Statistical significance was established by two-way ANOVA with Tukey’s multiple comparison test. ****p<0.0001 versus olaparib (50 mg/kg, QD). RBC: red blood cell. HGB: hemoglobin. RETIC: reticulocytes. PLT: platelet.

While PARPi have demonstrated clinical utility in HR deficient tumors, acquired resistance remains a significant therapeutic challenge^20^. Emerging evidences implicate BRCA reversion mutations mediated through MMEJ pathway as an important resistance mechanism of PARPi^7^. Based on this mechanistic understanding, we hypothesized that the combination of the Polθ inhibitor ZMS-6659 with PARPi might delay PARPi resistance development. To verify this hypothesis, we conducted a long-term study in an MDA-MB-436 tumor xenograft model, in which mice were daily oral-administrated with ZMS-6659 and olaparib as single agent or in combination for 78 days (Day 0-Day 77), followed by a 55-day treatment-free observation period (total duration of 133 days). Strikingly, tumors in the combination group showed complete regression (CR: 2/6 mice) at Day 77 and maintained durable suppression throughout the 55-day observation period after treatment cessation (CR: 3/6 mice). In contrast, olaparib treated tumors exhibited rapid recurrence following treatment withdrawal (Figure 4E). This sustained response pattern suggests that ZMS-6659 may disrupt the molecular pathways for PARPi resistance evolution.

Hematotoxicity remains a dose-limiting concern for PARPi monotherapy or combination strategies^21^. To systematically assess lineage-specific hemototoxicity risks, we developed a human CD34^+^ hematopoietic stem cell differentiation assay evaluating the toxicity on myeloid, erythroid, and megakaryocytic lineages. The result revealed that ZMS-6659 neither displayed significant effect on cell expansion and differentiation as a single agent nor enhanced the hematotoxicity in the combination with olaparib (Table 1). Consistent with these *in vitro* findings, the DLD-1 BRCA2-/- tumor xenograft model demonstrated that the ZMS-6659/olaparib combination therapy could elicit synergetic anti-tumor efficacy and result in tumor regression without enhancing the hematological toxicity (Figure 4F).

**Table 1.** The in vitro ZMS-6659 and olaparib combination hematotoxicity^a^.

| **Combo with olaparib** | **Myeloid** | **Erythroid** | **Megakaryocytic** |
| --- | --- | --- | --- |
| ZMS-6659 | >30000 | >30000 | >30000 |
| Olaparib alone | 4984 | 3578 | 393 |
| Olaparib combo with 10 μM ZMS-6659 | 7055 | 3240 | 379 |

a. Each data represents the mean from 3 biological replicates (n=3).

### Combination of ZMS-6659 and TOPOi shows synergetic antitumor effects

TOPOi can induce DSB accumulation in tumor cells and are widely used as chemotherapeutic agents in clinical treatment of tumors^15^. Given this clinical relevance, we hypothesized potential synergistic effects between TOPOi and Polθ inhibitors. We tested this hypothesis using two TOPOi, SN-38 (a Type I topoisomerase inhibitor) and etoposide (a Type II topoisomerase inhibitor). We found that, the Polθ expression was up-regulated after the treatment of both agents (Figure 5A & 5B), which was also found in HR deficient tumors^11^. Subsequent BrdU incorporation assays demonstrated enhanced suppression of DNA synthesis when combining ZMS-6659 with either SN-38 or etoposide, compared to single-agent treatments (Figure 5C & 5D), indicating a potentiated anti-proliferative activity through combination treatment. Then we further validated these results in the cell proliferation assay, and the combined treatment of ZMS-6659 and SN-38 or etoposide both elicit synergetic anti-proliferation activities in DLD-1 and NCI-H82, respectively (Figure 5E & 5F). Mechanistically, the combination therapy induced substantially elevated γH2AX levels (Figures 5G & 5H), accompanied by increased comet tail formation in single-cell electrophoresis assays (Figure 5I & 5J). These findings demonstrate that Polθ inhibition exacerbates TOPOi-induced genomic instability, resulting in enhanced cytotoxicity. Notably, this enhanced efficacy occurred in HR proficient contexts, suggesting that Polθ inhibition may provide therapeutic benefits beyond current biomarker-defined HR deficient populations.

**Figure 5.**
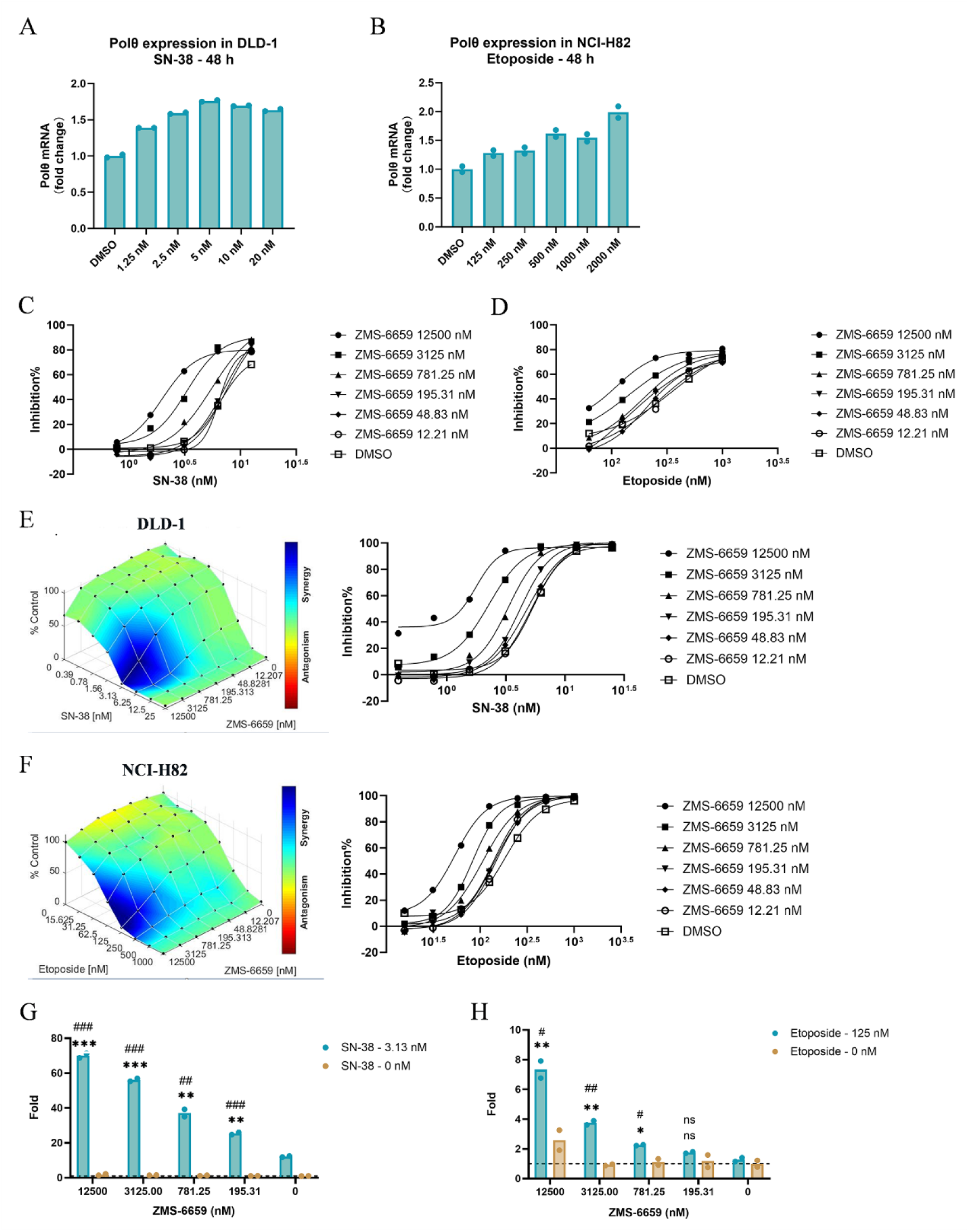
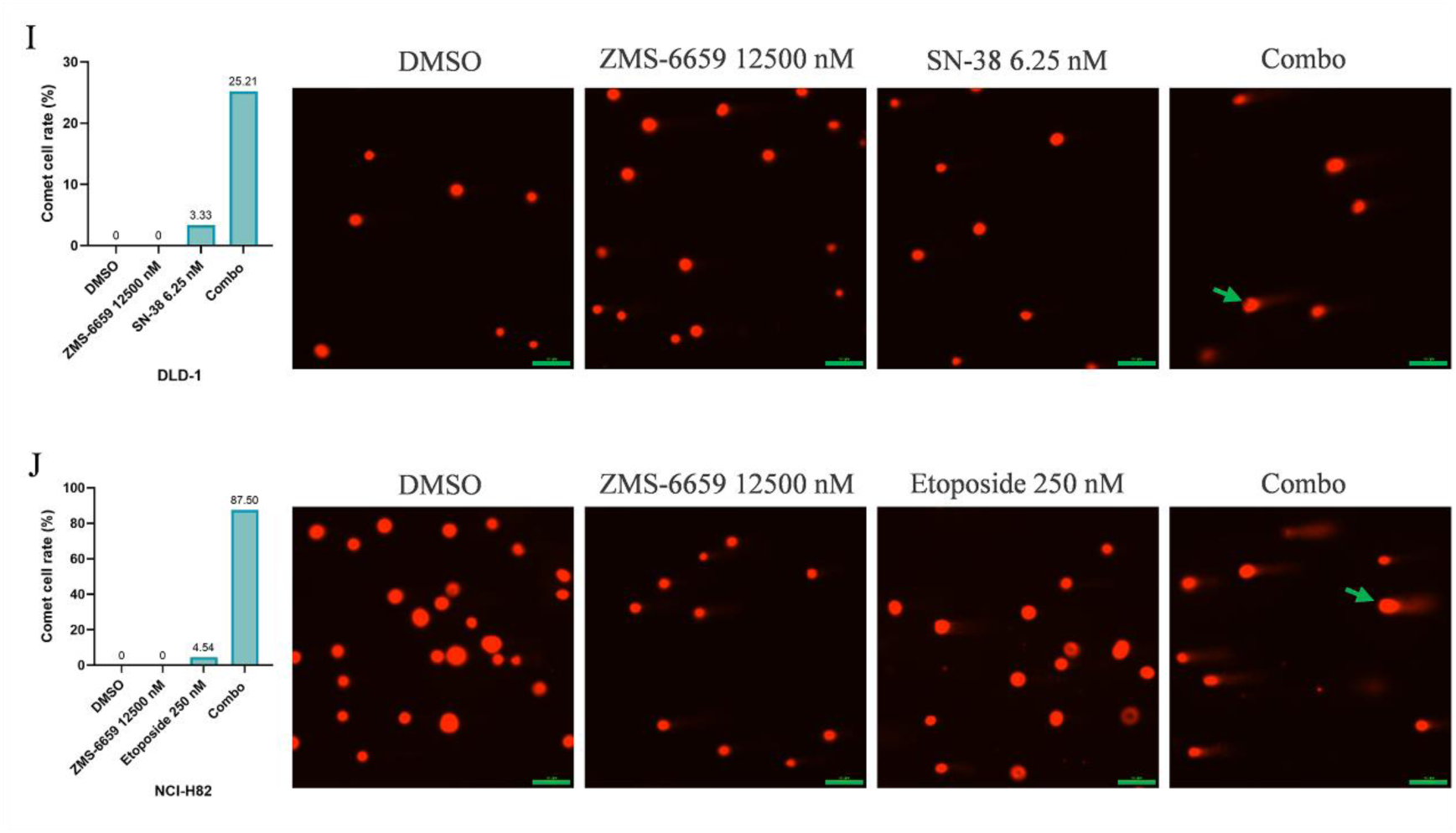
Combination of ZMS-6659 and TOPOi shows synergetic antitumor effects. (A & B) Polθ mRNA levels after the treatment of indicated concentrations of SN-38 (A) or Etoposide (B) for 48 h. Each data point represents the mean from 2 biological replicates (n=2). (C & D) BrdU tests for the ZMS-6659 and SN-38 combination (C) or the ZMS-6659 and Etoposide combination (D) inhibition of DNA synthesis. Each data point represents the mean from 2 biological replicates (n=2) (C) or from 3 biological replicates (n=3) (D). (E & F) The combination of ZMS-6659 and SN-38 in DLD-1 cells (E) or the combination of ZMS-6659 and Etoposide in NCI-H82 cells (F). Each data point represents the mean from 2 biological replicates (n=2). (G & H) γH2AX accumulation induced by the combination of ZMS-6659 with SN-38 in DLD-1 cells (G) or the combination of ZMS-6659 with Etoposide in NCI-H82 (H). Each data point represents the mean from 2 biological replicates (n=2). Statistical significance was established by unpaired t test: n.s. p≥0.05, *p<0.05, **p<0.01, ***p<0.001 versus SN38 or Etoposide. n.s. p≥0.05, #p<0.05, ##p<0.01, ###p<0.001 versus ZMS-6659 at the corresponding concentration. (I & J) Comet assays for DNA damage induced by the ZMS-6659 and SN-38 combination (I) or ZMS-6659 and Etoposide (J) combination. Each sample were imaged for 5 areas. Cells with the DNA tail were counted and bars stand for average ratio of cells with the DNA tail. The green lines were scales for 100 μM. The green arrows indicate example cells with the DNA tail.

## Discussion

Polθ is a unique DNA polymerase as it contains a helicase domain^4^. The helicase domain functions upstream of the polymerase domain and is responsible for the removal of RPA from ssDNA and ssDNA annealing, while the polymerase domain is required for the ssDNA extension. Our data demonstrate that the inhibition of either domain blocks Polθ-dependent MMEJ activity (Figure 2A). The first reported Polθ inhibitor ART558 targets the polymerase domain and exhibits moderate cellular activity in a nearly micromolar level^8^. A recent study identified RTx-161, a novel Polθ polymerase inhibitor, which exhibits improved biochemical potency, but still requires micromolar levels to inhibit HR deficient cell proliferation and elicit synergistic effects with PARPi^22^. Efforts to develop Polθ inhibitors by targeting the helicase domain have also been made. NVB has been demonstrated its ability to inhibit Polθ ATPase activity and is currently in Phase I clinical trial^17^. However, preclinical data indicates its weak biochemical activity with an IC_50_ of 24 μM, and its clinical efficacy remains unclear. Here, we report ZMS-6659, a novel Polθ helicase inhibitor, which displayed significantly stronger preclinical activities than previously reported Polθ polymerase or helicase inhibitors. It potently blocks proliferation of HR deficient DLD-1 BRCA2-/- cells with a digit nanomolar IC_50_, and shows over 900-fold selectivity over DLD-1 parent cells and non-malignant cells.

While ZMS-6659 exhibited synthetic lethality with HR deficiency *in vitro*, its limited anti-tumor efficacy as a monotherapy *in vivo* prompted us to investigate potential compensatory mechanisms. The shieldin complex (composed of SHLD1, SHLD2, SHLD3, and REV7) has been implicated in directing DSB repair toward NHEJ. In the situation of BRCA1 deficiency, it may be recruited by 53BP1 to the DSB sites to protect DNA ends from excessive end-resection, promoting DSB repair by NHEJ pathway and impairing Polθ inhibitor sensitivity^8,23–26^. To explore the effect of Shieldin complex on the sensitivity of ZMS-6659, we generated a SHLD2 knockout model in the natural BRCA1-mutated MDA-MB-436 cell line (MDA-MB-436 SHLD2-/-), and observed a dramatic improvement in the sensitivity to ZMS-6659 both *in vitro* and *in vivo* (Figure 2E-2G). This finding suggests therapeutic potential for ZMS-6659 as monotherapy in cancer patients concurrently harboring BRCA1 and shieldin complex deficiencies. Data from the cBioPortal database reveal that the concurrence of Shielding complex deficiency and BRCA1 mutation in solid tumors is about 1%, and it reaches 2%-6% in uterine endometrioid cancer, colorectal cancer and lung adenocarcinoma. Moreover, several studies have demonstrated that the deletion of any components in Shieldin complex may cause resistance to PARP inhibitor in BRCA1 mutant tumor cells due to the restoration of HR^8,23,24^. But we did not observe resistance to olaparib in MDA-MB-436 SHLD2-/-cell line (Figure S3), and suspected it may due to the different BRCA1 mutation patterns. MDA-MB-436 carries a BRCA1 5396+1G>A mutation that results in the loss of BRCA1 protein expression, which is different from cell lines carrying BRCA1 exon-11 mutation and expressing truncated BRCA1 variants. In such truncation mutants, shieldin deficiency enables residual BRCA1 to recruit PALB2 and facilitate RAD51-mediated HR restoration^27^.

Current clinical development of Polθ inhibitors predominantly focuses on combination strategies with PARPi, a paradigm supported by our findings demonstrating robust synergy between ZMS-6659 and PARPi. While PARPi monotherapy has demonstrated clinical efficacy in HR-deficient tumors, dose-limiting hematotoxicity remains a therapeutic challenge^21,28^. Existing strategies to enhance PARPi sensitivity through combination with chemotherapeutics or cell cycle checkpoint WEE1 inhibitors have been constrained by overlapping hematotoxicity profiles^29^, highlighting the needs for novel strategies with improved safety margins. To evaluate the hematopoietic toxicity of Polθ inhibitor in combination with olaparib, we developed a CD34^+^ differentiated lineage-specific hematotoxicity assay and observed that ZMS-6659 alone showed a low risk in hematotoxicity and did not enhance the hematotoxicity in combination with olaparib (Table 1). Hematotology analysis in the DLD-1 BRCA2-/- xenograft model further demonstrated no significant changes in red blood cells, platelets and hemoglobin in the combination group versus PARPi monotherapy, despite achieving robust synergy effect and inducing tumor regression (Figure 4F). This favorable safety profile may stem from Polθ’s restricted expression in normal tissues, with real-time PCR analysis showing 3-to 26-fold higher in tumor samples than in normal tissues^30^. Besides, dose-response exploration revealed that the reduced-dose of PARPi (15 mg/kg olaparib) combined with ZMS-6659 also achieve a significant increasing anti-tumor activity comparing with PARPi alone, demonstrating an opportunity to reduce PARPi dosage to improve the tolerance in clinical (Figure 4B). These findings collectively address a potential for mitigating PARPi on-target toxicities while maintaining therapeutic efficacy, and provide the mechanistic rationale for clinical prioritization of Polθ/PARPi combinations.

Beyond the combination with PARPi, the mechanistic rationale extends the Polθ combination strategies to other DNA-damaging therapies. Our results demonstrated that TOPOi induced compensatory up-regulation of Polθ expression, and ZMS-6659 in combination with topoisomerase inhibitors displayed a robust synergistic effect, accompanying with the accumulation of DNA damage (Figure 5). Notably, this synergy occurred in HR proficient cells, suggesting therapeutic potential beyond HR deficiency paradigms. A similar result was also reported in a study in which the combination of PARPi and IMMU-132, a trop2 ADC composed of a TOPOi payload, results in synergistic tumor growth inhibition and increased dsDNA breaks, regardless of BRCA1/2 status^31^. Since TOPOi display greatest cytotoxicity during S phase and induce S/G2 arrest^32–34^, and HR and MMEJ are the dominant repair pathways in the S/G2 phase, we speculated that the DNA damage induced by TOPOi might be too intense to be repaired by the HR repair pathway, and it is necessary to up-regulate Polθ to exert the repair function of MMEJ, thus generating a synergistic effect between TOPOi and Polθ inhibitor. A recent study also offers another possibility. The study identified Polθ-mediated MMEJ as a dominant repair pathway during mitosis, while both HR and NHEJ are repressed in mitosis^35^. We proposed that HR insufficiency during S/G2 phase leads to unresolved DSB, which subsequently requires Polθ-MMEJ intervention during mitosis. PARPi have been explored for the combination with TOPOi in clinical trials, and these combinations are very myelosuppressive and requires optimization of dosing or scheduling^32,36^. Our hematotoxicity profiling suggests Polθ inhibitors may circumvent this limitation, providing a potential advantage for the combination of Polθ inhibitors and TOPOi.

In this study, we reported a highly potent Polθ helicase inhibitor ZMS-6659 and its potential application as monotherapy or in combination with PARPi and TOPOi. Evidences suggest that MMEJ pathway in mitosis may account for the synthetic effect between Polθ inhibition and PARPi induced S phase DNA damage in BRCA2 mutant cells^35^. Our study did not figure out whether the synergy between Polθ inhibition and TOPOi is also related to the role of MMEJ in mitosis, and whether it distinguishes between different biomarkers. It is unclear that whether Polθ inhibition has potential to combine with mitotic inhibitors, such as anti-microtube agents. Clarifying these questions may help us to deeply understand the mechanism of Polθ inhibitor and its clinical positioning.

## Methods

### Enzymatic activity assay

Polθ (M1-N899) was synthesized by Pharmaron Inc.. WRN (522-1242) and BLM (641-1290) was synthesized by WuXi AppTec.. dT50 and Hec1 were synthesized by GenScript. ADP-Glo™ Kinase Assay was purchased from Promega. Compounds with series dilution (3-fold, 10 concentrations) were dispensed into 384-well plates by Echo 650 Liquid Handler. The final concentrations of compounds were started at 10 μM, 20 μM, 50 μM or 100 μM respectively, and the final concentration of DMSO was 0.6%. Enzymes, dT50, Hec1 and ATP were dispensed into plates that had been pre-dispensed with compound and incubated at room temperature. The final concentrations of Polθ (M1-N899), WRN (522-1242), BLM (641-1290), dT50, Hec1 were 1.5 nM, 1.5 nM, 2.5 nM, 10 nM and 0.2 nM separately. Luminescence signals (RLU) were detected by Envision plate reader. XLfit550 was used to analyze the data, and a four-parameter logistic regression model was used to fit the dose-response curve and calculate IC_50_.

The enzymatic activity assays of RecQ1, RecQ4, HELQ and FANCJ were performed by ICE Bioscience InC.. The enzymatic activity assays of SMARCA2 and SMARCA4 were performed by Pharmaron Inc..

For polymerase activity, Polθ (S1819-V2590) was synthesized by Pharmaron Inc.. The template and primer sequences were synthesized by Genewiz. dNTP was purchased from Sigma. Quant-iT PicoGreen dsDNA reagent was purchased from Invitrogen. The template and primer sequences were annealed to generate 10 μM primer-template duplex: 95°C for 5 min and ramp down to 25°C. Compounds with series dilution (3-fold, 10 concentrations) were dispensed into 384-well plates by Echo 650 Liquid Handler. The final concentrations of compounds were started at 6 μM, and the final concentration of DMSO was 1%. The enzyme, primer-template duplex and dNTP were dispensed into plates that had been pre-dispensed with compound and incubated at room temperature. The final concentrations of Polθ (S1819-V2590), primer-template duplex and dNTP were 2.5 nM, 25 nM and 20 μM separately. Fluorescence signals at the excitation/emission wavelengths of 485 nm/535 nm were detected by Envision plate reader. XLfit550 was used to analyze the data, and a four-parameter logistic regression model was used to fit the dose-response curve and calculate IC_50_.

### MMEJ reporter assay

The MMEJ reporter assay was performed by ICE Bioscience InC.. Before the experiment, the Neon transfection system was sterilized with UV light, and E2 and R buffer were warmed up to room temperature. HEK293T cells were cultured in DMEM containing 10% heat-inactivated FBS. Cells were trypsinized at confluence around 80-90% in T75 flask and suspended in culture medium. After counting, cells were resuspended in R buffer containing MMEJ substrate. Set Neon system, insert Neon tube into Neon Pipette Station, add E2 buffer into the tube, and start electroporation. The transfected cells were loaded into culture medium without antibiotics and seeded into 384-well plates. Cells were incubated at 37°C in a humidified CO_2_ incubator for 24 hours. The Nano-Glo reagent was pre-warmed to room temperature and mixed with luciferase substrate with a ratio of 50:1. Cell culture plates were equilibrated at room temperature for 10 min. 40 μL/well Nano-Glo was added into the plates, and the plates were shaked at 300 rpm for 3 min in the dark. Luminescence signals (RLU) were detected by BMG PHERAstar FSX. XLfit (5.5.0.5) was used to analyze the data, and a four-parameter logistic regression model was used to fit the dose-response curve and calculate IC_50_.

### Cell culture

The DLD-1 BRCA2-/- cells and DLD-1 parental cells were purchased from Horizon Discovery Inc.. MDA-MB-436 parental cells were purchased from Cobioer Biosciences Co., Ltd.. MDA-MB-436 SHLD2-/- cells were generated by Kyinno Co., Ltd.. NCI-H82 cells were purchased from ATCC. All cells were cultured with RPMI-1640 medium (Gibco) supplemented with 10% (v/v) fetal bovine serum (FBS). Cells were passaged at 80%-90% confluency and maintained at 37°C, 5% CO_2_.

### Cell proliferation

Cells were seeded into 96-well plates as follows: DLD-1 BRCA2-/- cells were seeded at the density of 1000 cells/well, DLD-1 parental cells were seeded at the density of 600 cells/well, MDA-MB-436 SHLD2-/- cells and MDA-MB-436 parental cells were seeded at the density of 3000 cells/well, NCI-H82 cells were seeded at the density of 1000 cells/well, HL-7702 cells were seeded at the density of 1000 cells/well. The plates were incubated at 37°C, 5% CO_2_ overnight. On the second day, compounds were diluted to required concentrations and added into cells respectively. The final concentration of DMSO was 0.25%. Cells were treated for 7 days at 37°C, 5% CO_2_. 50 µL/well CellTiter-Glo (CTG, Promega) reagent was added into cell culture plates and the plates were incubated at room temperature for 10 min. The luminescent signal values (RLU) were detected by using Envision plate reader.

XLfit550 was used to analyze the data, and a four-parameter logistic regression model was used to fit the dose-response curve and calculate IC_50_.

The cytotoxicity screening in human hepatocytes assay was performed by Pharmaron Inc.

### Flow cytometry for DNA damage marker γH2AX

Cells were seeded into 24-well plates as follows: DLD-1 BRCA2-/- cells were seeded at the density of 15000 cells/well, MDA-MB-436 SHLD2-/- cells were seeded at the density of 30000 cells/well. The cell plates were incubated at 37°C, 5% CO_2_ overnight. On the second day, compounds were diluted to required concentrations and added into cells respectively. The final concentration of DMSO was 0.2%. Cells were treated for 3 days at 37°C, 5% CO_2_. After 3-day treatment, cells were collected and transferred to a 96-well plate (V-bottom). Cells were washed with cell staining buffer twice and fixed with pre-cooled IC fixation buffer at 4°C for 40 min. Then, cells were washed with cell staining buffer twice and treated with pre-cooled True-PhosTM perm buffer at 4°C for 40 min. Next, Cells were washed with cell staining buffer twice and stained with Phospho-Histone H2A.X (Ser139) (20E3) Rabbit mAb (Pacific Blue™ Conjugate) at room temperature for 1 h. After washed with cell staining buffer twice, cells were stained with PI reagent (PI 30 μg/mL, RNase A 100 μg/mL, Triton X-100 0.2%) and flow cytometer was used to measure the percentage of cells with γH2AX signal. GraphPad Prism 10 was used to analyze the data.

### 3D bioprinting assay

3D bioprinting assay in patient-derived tissue specimen from one ovarian tumor was carried out by Cyberiad Life Sciences. The samples were cut into pieces and enzymatically digested by collagenase type I (Gibco, 17018029). The resulting cell suspension was mixed with ovarian bioink (Cyberiad, YJP0102) and 3D ovarian models constructs were fabricated using a 96-well 3D bioprinter (Cyberiad, Biocube) at a density of 20000 cells per well. 3D ovarian models were incubated overnight at 37 °C, 5% CO₂. Compounds were diluted to required concentrations and added on the next day. Constructs were incubated for 9 days at 37 °C, 5% CO₂, with medium changes performed every 2–3 days. Cell viability was assessed using the CellTiter-Glo 3D assay (Promega, G9683), and luminescence was measured with a plate reader (Tecan, Infinite E Plex).

### Animals

All experimental procedures involving animals and their care were conducted in conformity with the State Council Regulations for Laboratory Animal Management (Enacted in 1988) and were approved by the Institutional Animal Care and Use Committee of Simcere Biology Medical Technology Co., Ltd., People’s Republic of China. Female NU/NU mice at 6–8 weeks of age were purchased from Beijing Vital River Laboratory Animal Technology Co., Ltd.. Female NPG mice at 7–8 weeks of age were purchased from Beijing Vitalstar Biotechnology Co.,Ltd.. Female NOD SCID mice at 7–8 weeks of age were purchased from Shanghai Lingchang Biotechnology Co., Ltd..

### Xenograft models

Female NU/NU mice were subcutaneously injected with 100 μL 1 × 10^7^ DLD-1 BRCA2-/- cells into the right flank. Female NOD SCID or NPG mice were subcutaneously injected with 100 μL 1 × 10^7^ MDA-MB-436 parental cells or MDA-MB-436 SHLD2-/- cells into the right flank, respectively. Mice were randomized into different groups with mean tumor volume around 100 mm^3^. Mice were then received daily administration of vehicle, ZMS-6659, Olaparib or ZMS-6659 and Olaparib as indicated. Body weights and tumor volumes were measured twice a week. Blood samples were collected at indicated time points for PK and blood sample tests.

### In vitro hematotoxicity

The in vitro human CD34^+^ hematopoietic stem cell differentiation assay was developed according to a pre-described reference^16^. Frozen vials of mobilized peripheral stem/progenitor CD34^+^ cells were rapidly thawed in a 37°C water bath. Cells were transferred to 10 mL HBSS with 10% FBS and centrifuged for 5 min at 300 × g. The supernatant was removed and cells were re-suspended in 2 mL StemSpan SFEMII basal medium. Cells were seeded into ultra-low-attachment, round-bottom 96-well plates at the density of 2000 cells/well with lineage-specific cytokine combinations: myeloid with 1% CC100 cytokine cocktail (100×) and 20 ng/mL GM-CSF, erythroid with 1% Erythroid supplement (100×), megakaryocytic with 1% Megakaryocyte supplement (100×). Compounds were diluted to required concentrations and added into cells respectively. The final concentration of DMSO was 0.3%. Cells were treated for 14 days at 37℃, 5% CO_2_. The plates were gently shaken daily (for 2 to 3 min). 50 µL/well CellTiter-Glo 3D (CTG 3D, Promega) reagent was added into cell culture plates and the plates were incubated at room temperature for 30 min. The luminescent signal values (RLU) were detected by using Envision plate reader. XLfit550 was used to analyze the data, and a four-parameter logistic regression model was used to fit the dose-response curve and calculate IC_50_.

### Quantitative PCR (qPCR/qRT-PCR)

Quantitative PCR assay was performed according to FastPure Cell/Tissue Total RNA Isolation Kit V2 and HiScript II One Step qRT-PCR SYBR Green Kit (Vazyme). Cells were collected and lysed by 500 μL Buffer RL 1. The lysate was transferred to FastPure gDNA-Filter Columns III and centrifuged at 13400 × g for 1 min. The filtrate was collected, and absolute ethanol (0.5 × the volume of filtrate) was added and mixed with the filtrate thoroughly. The mixture was transferred to FastPure RNA Columns III and centrifuged at 13400 × g for 1 min. The filtrate was discarded, and 700 μL Buffer RW1 was added to the FastPure RNA Columns III and centrifuged at 13400 × g for 1 min. The filtrate was discarded, and 700 μL Buffer RW2 was added to the FastPure RNA Columns III and centrifuged at 13400 × g for 1 min. The filtrate was discarded, and 500 μL Buffer RW2 was added to the FastPure RNA Columns III and centrifuged at 13400 × g for 5 min. The filtrate was discarded, and the FastPure RNA Columns III was centrifuged at 13400 × g for 2 min and placed at room temperature for 10 min. 50 μL of RNase-free ddH_2_O was added to the adsorption column, and the adsorption column was incubated at room temperature for 5 min and centrifuged at 13400 × g for 1 min to elute the RNA. The amount of total mRNA transcripts was measured with NanoDrop 8000 (Thermo Fisher Scientific). The qPCR reaction system and program were set according to the HiScript II One Step qRT-PCR SYBR Green Kit. The qPCR program was run with QuantStudio™ 6 (Thermo Fisher Scientific). GraphPad Prism 10 was used to analyze the data.

### BrdU ELISA assay

BrdU ELISA assay was performed according to Cell Proliferation ELISA, BrdU (chemiluminescent) assay kit (Roche). Cells were seeded into 384-well plates as follows: DLD-1 parental cells were seeded at the density of 1500 cells/well, NCI-H82 cells were seeded at the density of 2000 cells/well. The plates were incubated at 37°C, 5% CO_2_ overnight. On the second day, compounds were diluted to required concentrations and added into cells respectively. The final concentration of DMSO was 0.25%. Cells were treated for 48 h at 37°C, 5% CO_2_ and labeled with BrdU (4 uL/well, 1:100 dilution) for 2 h at 37°C. The supernatant was removed and cells were incubated with FixDenat solution (80 uL/well) for 30 min at 25°C. FixDenat solution was removed and cells were incubated with Anti-BrdU POD (40 uL/well, 1:100 dilution) for 1.5 h at 25°C. Antibody conjugate was removed and cells were rinsed three times with Washing solution. The supernatant was removed and cells were incubated with Substrate Solution (40 ul/well, 1:100 dilution) for 5 min at 25°C. The luminescent signal values (RLU) were detected by using Envision plate reader.

### Comet assay

Comet assay was performed according to Comet Assay Kit (Beyotime). 1% normal melting point agarose was prepared and added onto the Comet Assay Slide (Beyotime) to create a base layer. Cells were collected and mixed with 0.7% low melting point agarose, and the mixture was added onto the base layer. The slide was transferred to pre-chilled lysis buffer and incubated at 4°C overnight. After cell lysed, the slide was rinsed with PBS (Beyotime) 3 times. The slide was transferred to a horizontal electrophoresis chamber containing pre-chilled alkaline solution (1 mM EDTA (pH8.0), 200 mM NaOH, pH 13) and incubated at room temperature for 30 min. Apply voltage to the chamber for 25 min at 25 volt. Then, the slide was transferred to pre-chilled neutral solution (0.4M Tris-HCl, pH7.5) and incubated at 4°C for 10 min. 20 µL Propidium Iodide Solution was added onto the slide and incubated in dark for 20 min. The slide was covered with cover glass and viewed by epifluorescence microscopy using a FITC filter.

## Data Availability

The data generated in this study are available upon request from the corresponding author.

## ACKNOWLEDGMENT

We acknowledge specialized software including GraphPad Prism, Combenefit, and XLfit to analyze the data and create figures. Simcere Zaiming Pharmaceutical Co., Ltd. provided financial support to the current study.

## AUTHOR CONTRIBUTIONS

All authors read and approved the final version of the manuscript. *F. Z.*, *G. L.* and *L.T.X.*: contributed to conceptualization, validation, supervision, investigation, and writing-review and editing. *L.J.*, *J.J.L.*, *L.L.* and *M.Y.L*: contributed equally to conceptualization, methodology, analysis, and writing-original draft for this work. *W.J.L.*, *Z. L., S.Q.*, *M.Y.S.*, *J. D.*, *C.X.A.*, *Y.Q.*: contributed to investigation, methodology and data curation. *Z.T.L.*, *L.Z.*, *C.Y.*, *X.Y.W.* and *R.H. T.*: contributed to conceptualization, supervision and resources.

## Supplementary Information

**Supplementary Figure 1.**
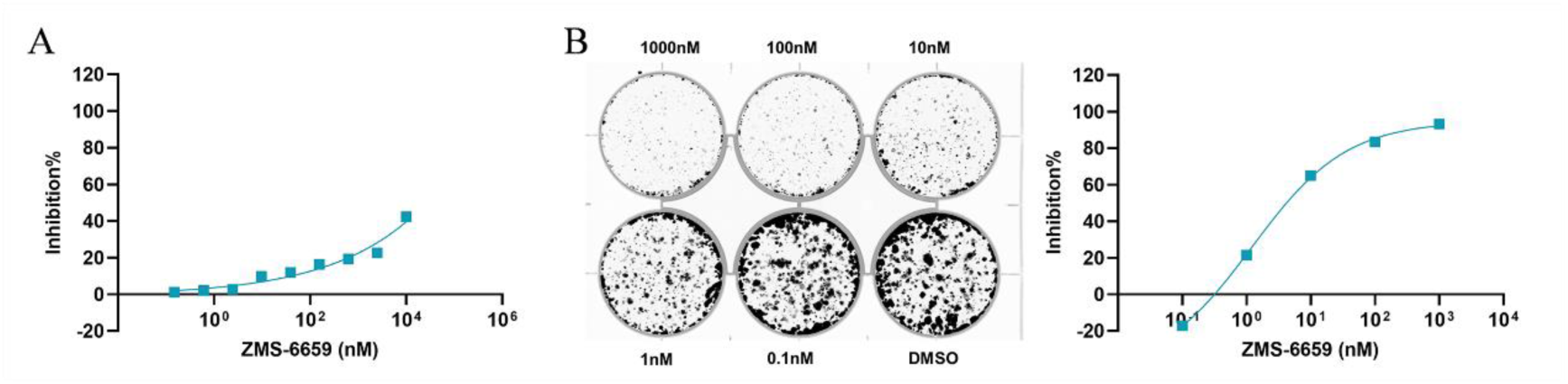
The anti-proliferation activity of ZMS-6659 in MDA-MB-436 cells. (A) The 7-day cell proliferation assay. Each data point represents the mean from 2 biological replicates (n=2). (B) The 14-day colony formation assay. The curve (right) was generated from the quantification of the original image (left).

**Supplementary Figure 2.**
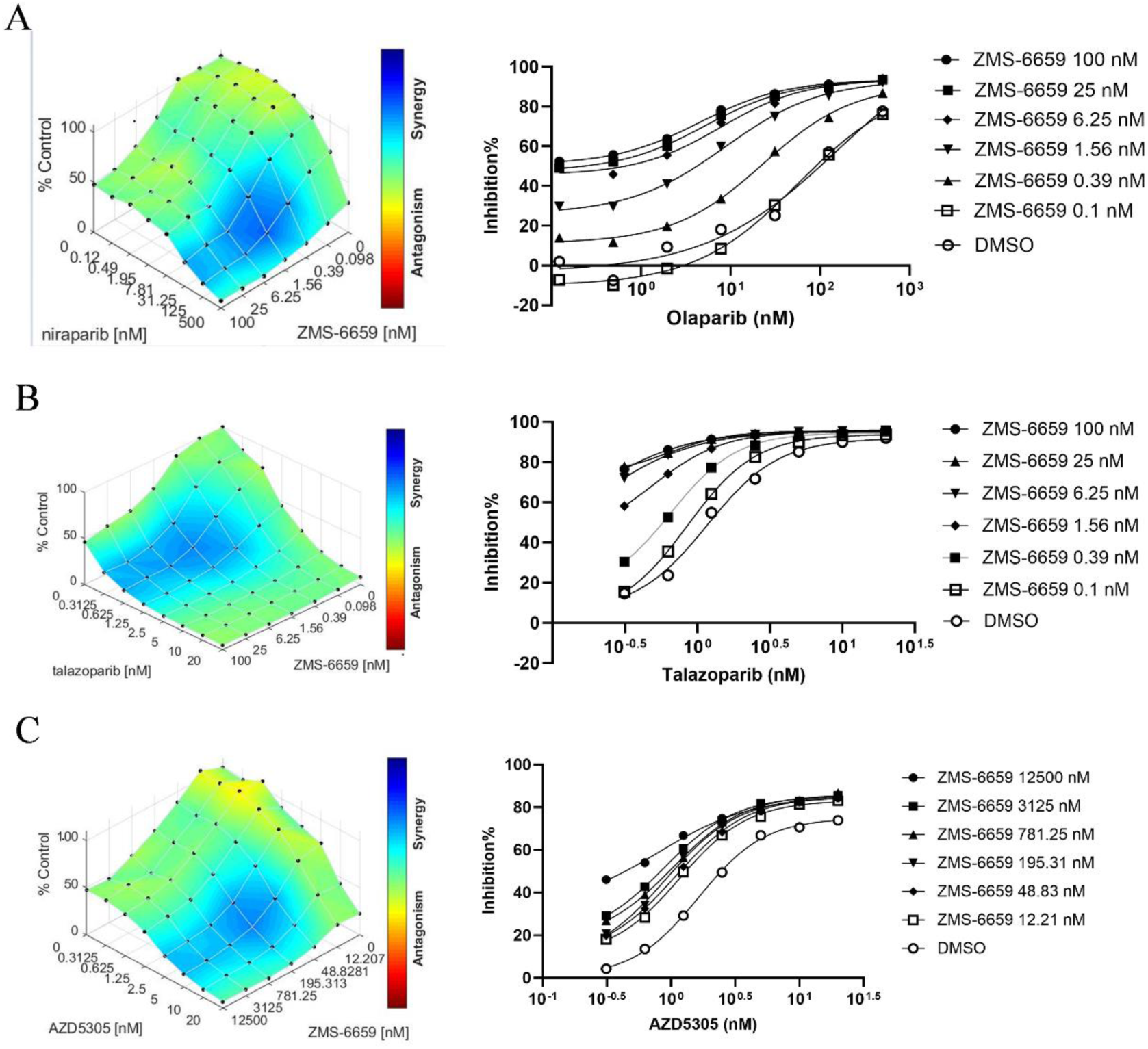
ZMS-6659 elicits synergy effect with PARPi. (A-B) The combination of ZMS-6659 and niraparib (A) or talazoparib (B) in DLD-1 BRCA2 -/- cells. (C) The combination of ZMS-6659 and PARP1-selective inhibitor AZD-5305 in MDA-MB-436 cells. Each data point represents the mean from 2 biological replicates (n=2).

**Supplementary Figure 3.**
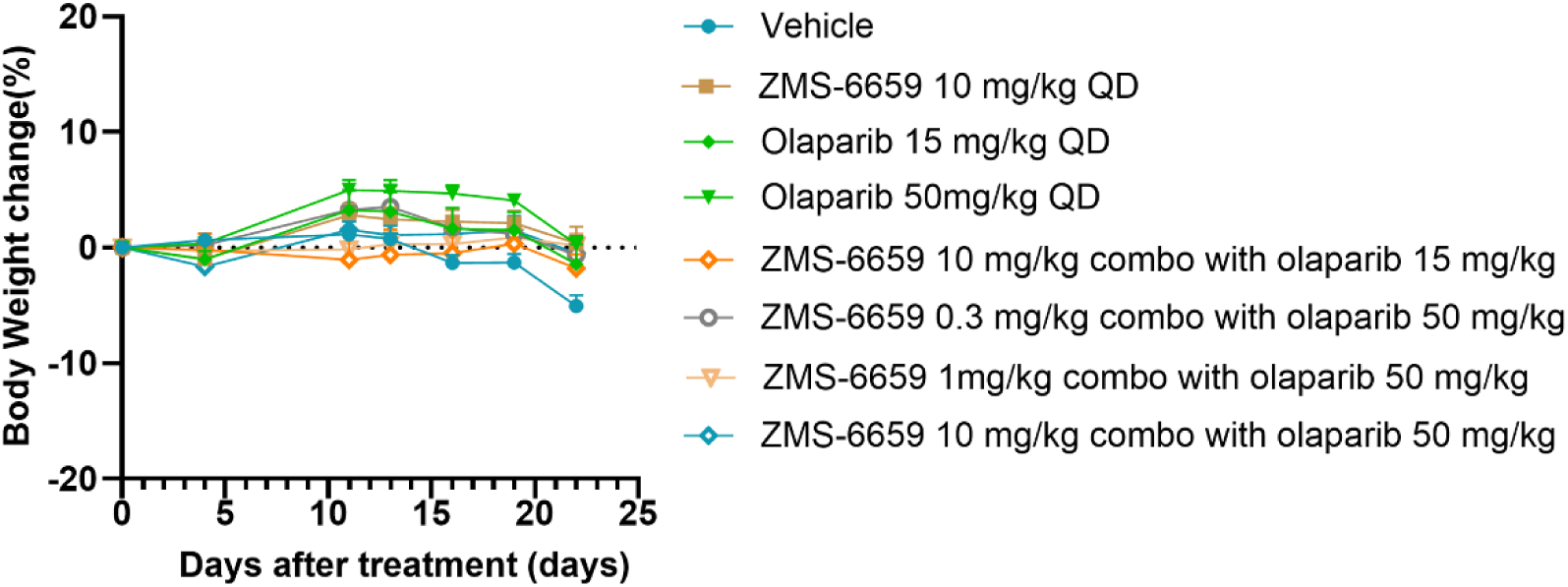
Mice body weight in the study of ZMS-6659 and olaparib combination treatment in DLD-1 BRCA2-/- xenograft mice model. Animals were treated for 22 days with either vehicle (n=8), or ZMS-6659 (10 mg/kg, QD, n=8), or olaparib (15/50 mg/kg, QD, n=8), or ZMS-6659 (10 mg/kg, QD) and olaparib (15 mg/kg, QD) combination (n=8), or ZMS-6659 (0.3/1/10 mg/kg, QD) and olaparib (50 mg/kg, QD) combination (n=8).

**Supplementary Figure 4.**
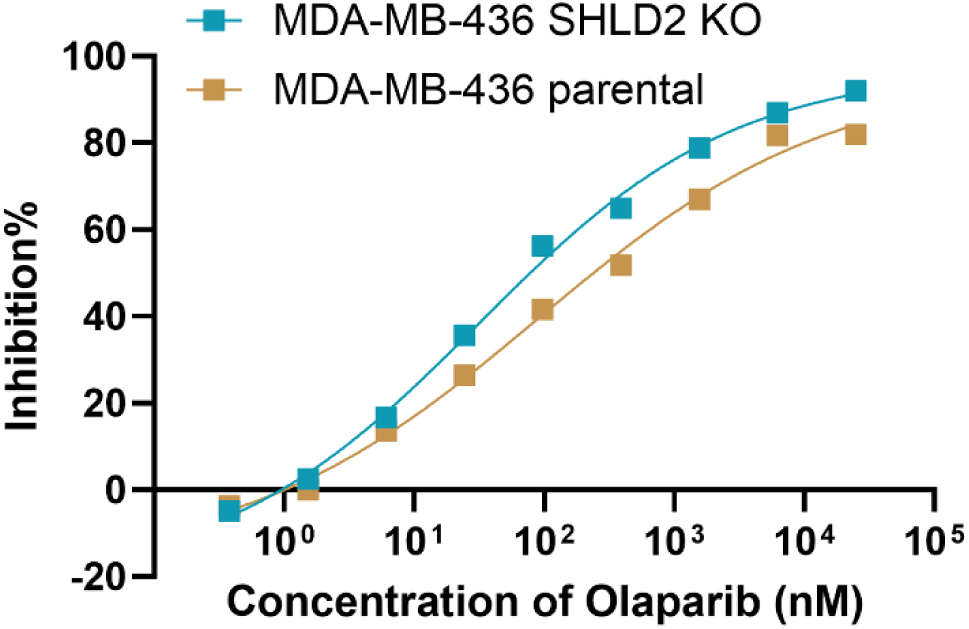
Cell proliferation inhibition activities of Olaparib in MDA-MB-436 SHLD2-/- and MDA-MB-436 parental cells. Each data point represents the mean from 2 biological replicates (n=2).

**Supplementary Table 1.**
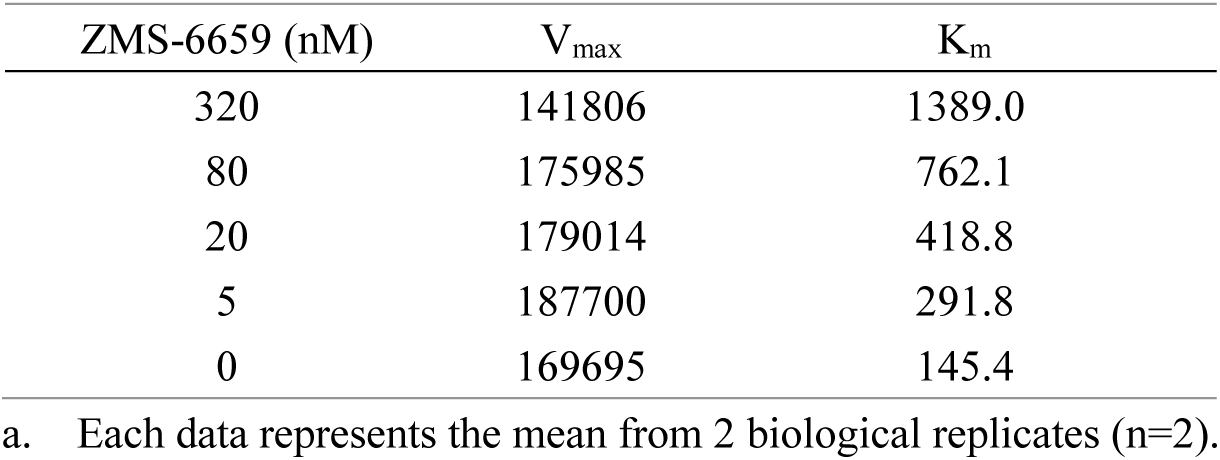
The V_max_ and K_m_ in the ATP competition assay^a^.

| ZMS-6659 (nM) | $V_{\max}$ | $K_m$ |
| --- | --- | --- |
| 320 | 141806 | 1389.0 |
| 80 | 175985 | 762.1 |
| 20 | 179014 | 418.8 |
| 5 | 187700 | 291.8 |
| 0 | 169695 | 145.4 |

a. Each data represents the mean from 2 biological replicates (n=2).

